# Dynamics, signals and influencing factors of CD4 T cell activation revealed by single cell RNA-seq

**DOI:** 10.1101/2022.08.13.503873

**Authors:** Hongyi Liu, Hui Li, Yifei Liu, Xuefei Wang, Shiya Yu, Xiangru Shen, Qi Zhang, Ni Hong, Wenfei Jin

## Abstract

T cell activation is a key event in adaptive immunity. However, factors affecting T cell activation have not been systematically analyzed. Here, we analyzed stimulated CD4 T cells with anti-CD3/CD28 under several conditions to explore the factors affecting T cell activation. We defined stimulated T overlapped with resting T on UMAP as inert T. Inert T expressed activated T specific genes and cytokines, indicating it is a special functional state. Stimulated T derived from peripheral CD4 T has higher fraction of effector T (T_EFF_) while stimulated T derived from CD4 T_N_ has higher fraction of proliferation T and interferon highly expressed T (IFN^hi^ T). CD4 T was more likely to differentiate into T_EFF_ and less likely to differentiate into heat shock protein specific T (HSP^hi^ T) and IFN^hi^ T in the presence of CD8 T. Interestingly, *CXCR4*^low^ T responded to stimulation more efficiently than *CXCR4*^hi^ T. These information facilitates we design stimulation to obtain ideal activated T.

## Introduction

CD4 T cells, which are a key component of adaptive immunity, not only play important role in elimination of pathogens, but also play important role in regulation of autoimmune disease and clearance of pathogenic cells such as cancer cells (*1-6*). If T cell receptor (TCR) on the surface of T cells recognizes an antigen-loaded MHC, the T cell will quickly differentiate and proliferate into a large number of effector T cells (T_EFF_) that recognize the same antigen, which is called T cell activation (*7-9*). T cell activation plays a critical role in establishing and controlling adaptive immune responses against pathogens and pathogenic cells, which is the central to adaptive immune response (*10, 11*). The most commonly used model to study CD4 T cell activation is the stimulation of naïve CD4 T cells (T_N_) with anti-CD3/CD28 beads. CD4 T_N_, which egress from thymus before they encounter any antigen, are fairly quiescent and can differentiate into different subtypes of T_EFF_ after T cell activation. Studies of T cell activation based on CD4 T_N_ have significantly increased our understanding of T cell activation process and cytokine-induced polarization (*12-15*). There are many studies analyzed the effects of cytokine sets on the differentiated T_EFF_ subsets. E.g., naïve CD4 T cells (T_N_) could be polarized by IFN-γ, IL-4, and IL-17 into Th1, Th2 and Th17, respectively (*16-19*).

Different from relatively homogeneous CD4 T_N_, CD4 T cells in human peripheral blood are a heterogeneous cell population including several T cell subsets including T_N_, central memory T cells (T_CM_), effector memory T cells (T_EM_), regulatory T cells (Treg) (*4, 20-22*). Different CD4 T cell subsets have different responses to stimulation. E.g. T_CM_ and T_EM_ quickly proliferate and secrete cytokines upon stimulation, while T_N_ responds to stimulation much slow and could differentiates into various T cell subsets under different microenvironment (*8, 23-25*). The development of single cell RNA-sequencing (scRNA-seq) provide a unique opportunity to investigate the effect of multiple factors on T cell activation. Recently, Cano-Gamez et al. (*20*) analyzed CD4 T_M_ and showed T cell activation are influenced by the effectorness gradient. CD4 T_N_ differentiates into Th2 with IL-4 polarization, while CD4 T_M_ could not differentiate into Th2 phenotype. Soskic, et al. identified lots of genetic variants that regulate gene expression dynamics during CD4 T cell activation (*26*). However, many factors affecting T cell activation have not been explored. In particular, there is no systematic study on how cellular heterogeneity and T cell communications affect on T cell activation.

Here, we stimulated peripheral CD4 T cells with anti-CD3/CD28 beads and performed scRNA-seq. Integrated analyses of CD4 T cells pre-and post stimulation identified CD4 T cell subsets such as T_N_, T_CM_, T_EM_, Treg, effector memory T cells re-expressing CD45RA (T_EMRA_), 4 different T_EFF_ subsets and HSP^hi^ T. We identified two main trajectories of T cell activation both of which are from T_N_ to T_EFF_, with differences lying in whether they pass through T_M_. We called the stimulated T cells overlapped with resting T cells on UMAP as inert T. Comparing with resting T, inert T highly expressed activated T specific markers. We conducted a series of comparison analyses on dynamics and signals of CD4 T cell subsets under various stimulations by integrated multiple datasets. We found that CD4 T cell subsets, cell-cell communications, and cytokines reprogrammed and reshaped activated T cell subsets, expressed genes and cell states.

## Results

### The T cell subsets pre-and post CD4+ T cell activation

We designed a workflow to analyze the dynamics/signals of T cell subsets and factors affecting T cell activation based on anti-CD3/CD28 stimulation of CD4 T cells (Fig. 1A). In brief, we isolated peripheral CD4 T cells by CD4 MicroBeads from a healthy donor. The peripheral CD4 T cells were stimulated with anti-CD3/CD28 magnetic beads (Dynabeads™ Human T-Activator CD3/CD28 for T Cell Expansion and Activation) for 16 hours. We performed single cell RNA-seq (scRNA-seq) on both the resting peripheral CD4 T cells and stimulated CD4 T cells using 10x Genomics Chromium™ Single Cell 3’ Library & Gel Bead Kit v2. After quality control, total 31,815 CD4 T cells, including 21,573 resting CD4 T cells and 10,242 stimulated CD4 T cells were left for further analyses. Uniform Manifold Approximation and Projection (UMAP) plot showed that resting CD4 T cells and stimulated CD4 T cells displayed quite different distribution (Fig. 1B). Resting CD4 T cells concentrated at the bottom, while stimulated CD4 T cells were widely distributed, with majority in the upper part (Fig. 1B). The expression level of active T cell marker *IL2RA (27*) and naïve T cell marker *KLF2* (*28*) was up regulated and down regulated, respectively (Fig. 1C). We identified 69 resting T cell specific genes and 253 stimulated T cell specific genes (fig. S1, A and B), majority of which consist with our previous study by bulk data (*29*).

**Fig. 1.**
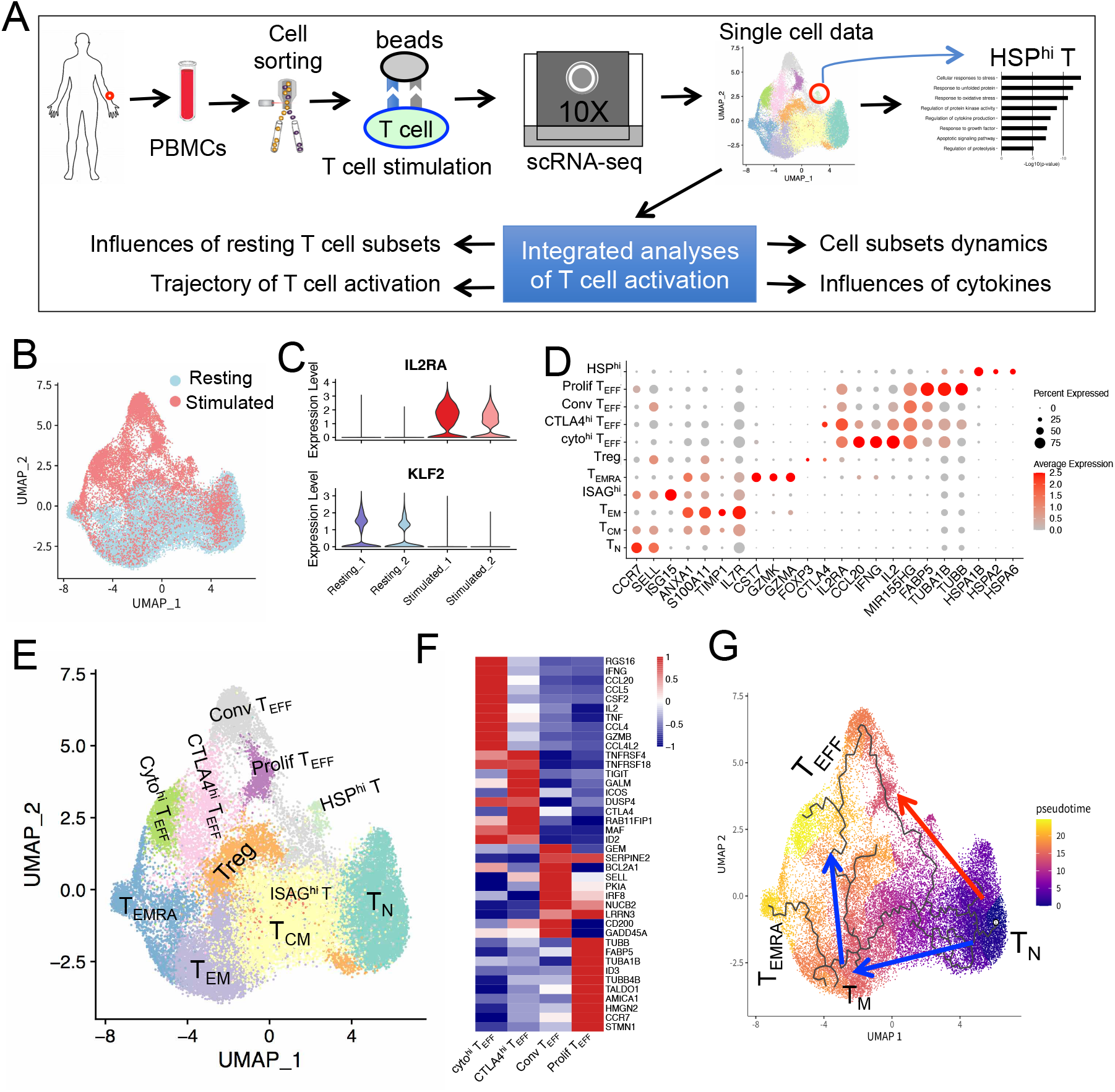
Scheme of this study and T cell subsets pre-and post-stimulation. **(A)** Scheme of this study. **(B)** UMAP projection of CD4 T cells pre-and post-anti-CD3/CD28 stimulation, colored by whether it was stimulated. **(C)** Expression level of *IL2RA* and *KLF2* in resting T cells and stimulated T cells. IL2RA is T cell activation markers, while KLF2 is naïve T cell marker. **(D)** Normalized expression level and expression percentage of cell-type-specific genes in 11 T cell subsets. **(E)** UMAP projection of CD4 T cells pre-and post-anti-CD3/CD28 stimulation, colored by cell subsets. **(F)** Heatmap of expression level of DEGs between conv T_EFF_, CTLAA4^hi^ T_EFF_, cyto^hi^ T_EFF_ and prolif T_EFF_. **(G)** Two major T cell activation trajectories inferred by Monocle 3, colored by cell’s pseudotime value.

We identified 11 T cell subsets and annotated them according to their specific expressed marker genes (Fig. 1, D and E). There are 6 T cell subsets mainly from resting CD4 T cells, namely T_N_ (*CCR7*^hi^, *SELL*^hi^), T_CM_ (*CCR7*^+^, *SELL*^+^, *ANXA1*^+^), T_EM_ (*CCR7*^*low*^, ANXA1^+^, *TIMP1*^*+*^), CD4 T_EMRA_ (*ANXA1*^+^, *CST7*^*+*^, *GZMK*^*+*^, *GZMA*^*+*^), Treg (*FOXP3*^+^, *CTLA4*^+^) and IFN signaling associated gene (ISAG) highly expressed T cells (ISAG^hi^ T) (*ISG15*^hi^, *CCR7*^*+*^, *SELL*^+^) (Fig 1, D and E). These T cell subsets are essentially consistent with recent studies on peripheral CD4 T cells (*4, 20, 26*). There are 5 T cell subsets mainly from stimulated CD4 T cells, four of which belong to CD4 T_EFF_ (*IL2RA*^+^, *NME1*^+^ and *MIR155HG*^+^, *FABP5*^+^) (Fig 1D, E). There is a small T cell subset specifically expresses the heat shock protein (HSP) genes such as *HSPB1, HSPA1A* and *HSPA6* (Data S1) (*30*), which was called as HSP^hi^ T. We further compared the four CD4 T_EFF_ subsets, which is the dominant T cell subsets after anti-CD3/CD28 beads stimulation. We found each of the four T_EFF_ subsets has different feature and were called them as conventional T_EFF_ (conv T_EFF_), cytokines high T_EFF_ (cyto^hi^) T_EFF_, proliferation T_EFF_ (prolif T_EFF_) and CTLA4^hi^ T_EFF_ (Fig. 1D, 1F and fig. S1,C,D).

We inferred the pseudotime trajectories of these T cells for better understanding T cell activation processes (Fig. 1G). The inferred pseudotime contained two major trajectories: 1) T_N_ → T_EFF_; 2) T_N_ → T_CM_ → T_EM_ → T_EFF_. Both trajectories started from T_N_ and ended at T_EFF_, with different processes. The first trajectory initiated at T_N_ and directly differentiated into T_EFF_, which is short and the most direct T cell activation process. The second trajectory has one main path with two differentiation phases, from which derived several small branches. Phase #1 of the second trajectory is development of T_M_ from T_N_, which has two derived branches: 1) T_N_ → T_CM_ → T_EM_ → T_EMRA_; 2) T_N_ → T_CM_ → T_EM_ → Treg. The phase #2 of the second trajectory is the differentiated into T_EFF_ from T_M_ (Fig. 1G). The expressions of T cell activation markers (*MIR155HG, IER3*) and cytotoxicity related genes (*NKG7, GZMH*) were upregulated alongside lineage development (fig. S1E).

### Feature of HSP^hi^ T and HSP genes are major contributor for the formation of HSP^hi^ T cluster

HSP^hi^ T is a small T cell subset specifically expresses HSP genes such as *HSPA6, HSPA1A*, and *HSPA1B* (Fig. 2A-2B). HSP^hi^ T were solely from stimulated CD4 T cells, without contribution from resting T cells (Fig. 2C), indicating HSP^hi^ T is a specific state of stimulated T. Comparing with other T cells, HSP^hi^ T specifically expressed *HSPA1B, HSPA1A, HSPA2, HSPA6, METTL12, GADD45B, IER2, IER* and *DNAJB1* (Fig. 2D). These HSP^hi^ T specific genes enriched GO terms were cellular responses to stress (P=1.3×10^−13^), response to unfolded protein (P=2.5×10^−12^), response to oxidative stress (P=1.7×10^−11^), regulation of protein kinase activity (P=1.0×10^−9^), regulation of cytokine production (P=1.1×10^−8^), apoptotic signaling pathway (P=6.4×10^−8^) and regulation of proteolysis (P=6.2×10^−6^) (Fig. 2E). Based on the HSP^hi^ T specific enriched GO, HSP^hi^ T might be derived from T cells that over-response to anti-CD3/CD28 beads stimulation.

**Fig. 2.**
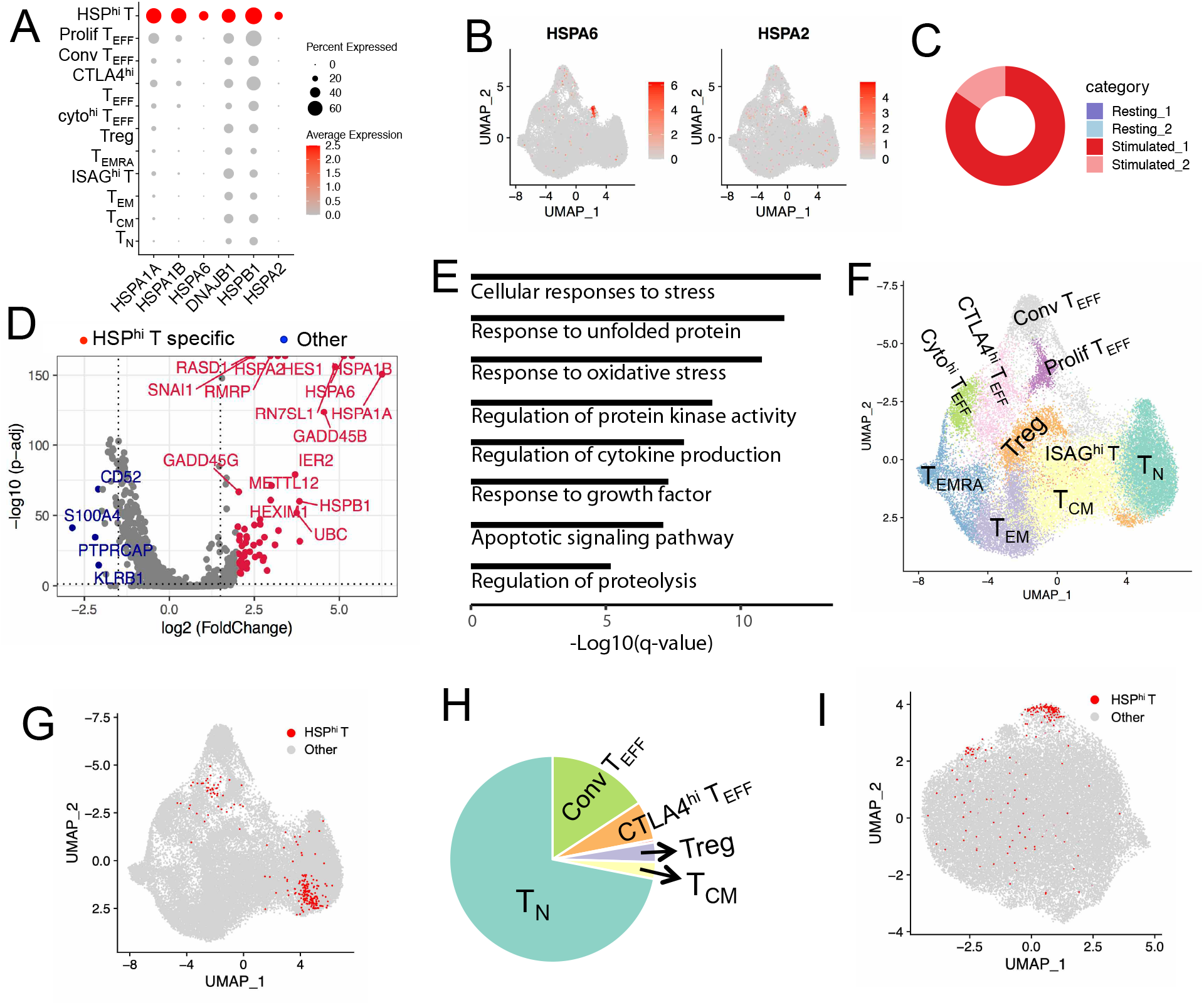
Feature of HSP^hi^ T. **(A)** Dot plot of HSP^hi^ T specific expressed genes. Color represents normalized expression level, and dot size represents expression percentage. **(B)** Expression level of *HSPA6* and *HSPA2* on UMAP plot of CD4 T cells. **(C)** Cell composition of HSP^hi^ T based on cell sources. **(D)** Volcano plot of DEGs between HSP^hi^ T and the other T cells (p < 0.01). Red points represent HSP^hi^ T cell specific high expressed genes, while blue points represent the other T cell specific highly expressed genes. **(E)** GO enrichment analysis of HSP^hi^ T specific expressed genes. **(F)** UMAP projection of CD4 T cells using gene sets that removed HSP related genes, in which HSP^hi^ T cell cluster disappears. **(G)** UMAP projection of CD4 T cells using gene sets that removed HSP related genes, with red points highlighting the original identified HSP^hi^ T cells. **(H)** Pie plot of number of HSP^hi^ T projected on each CD4 T cell subset, with 72% projected on T_N_. **(I)** UMAP projection CD4 T cells based on HSP related genes, with red points highlighting original identified HSP^hi^ T cells.

We re-conducted dimension reduction and clustering analysis on T cells using gene set excluded HSP genes. The re-clustered T cell subsets were the same as that using genome-wide data, except disappearance of HSP^hi^ T cluster. (Fig. 2F). In the re-projected UMAP, HSP^hi^ T was mainly projected on T_N_ (71.93 %), with moderate on conv T_EFF_ (15.78%) and a small fraction on CTLA4^hi^ T_EFF_ (6.14%), Treg (3.1%) and cyto^hi^ T_EFF_ (0.43%) (Fig. 2, G and H). The fraction of HSP^hi^ T compared to CD4 T_N_ is also the highest among all T cell subsets after we normalized the number of HSP^hi^ T compared to its projected T cell subsets (fig. S1F). Most HSP^hi^ T projecting on T_N_ indicated that gene expression profile of HSP^hi^ T has the highest similarity to that of CD4 TN if HSP genes were ignored. These results further indicated that majority of HSP^hi^ T was derived from CD4 TN that respond to stimulation of anti-CD3/CD28 beads. Interestingly, HSP^hi^ T cluster showed up when we only used HSP related genes to conduct UMAP (Fig. 2I), further indicating that HSP related genes were the major contributor for the formation of HSP^hi^ T cluster.

### Inert T and CXCR4 affecting T cell activation

The anti-CD3/CD28 stimulated T cells in the upper of UMAP plot are stimulation specific T cell subsets such as conv T_EFF_, cyto^hi^ T_EFF_, prolif T_EFF_ and HSP^hi^ T, thus we called them as activated T cells (Fig. 3A). The stimulated T cells that overlapped with the resting T cells on UMAP plot should have similar expression profile to that of resting T cells, thus we called them inert T cells (Fig. 3A). In this way, the stimulated T cells were separated into 6,316 activated T cells (61.67%) and 3,926 inert T cells (38.33%) (fig. S2A). Compared with resting T cells, inert T cells highly expressed cytokine related genes (*CCL3, CCL4, CCL3L3, CCL4L2, IL22, IFNG* and *STAT1*), and activated T specific genes (*IL2RA, NME1, MIR155HG, FABP5, IL2, CD200* and *IER3*) that are also highly expressed in activated T cells (Fig. 3B; Data S2). The most significantly enriched GO terms are cytokine signaling (P=2.09×10^−25^), response to stimulus (P=3.16×10^−22^), NF-κB signaling pathway (P=5.50×10^−17^) (Fig. S2B). These results indicate inert T cells are not completely quiescent and they are activated in someway if we only check the activate T cell markers. Compared with activated T cells, inert T cells highly expressed cytokine related genes (*CCL3, CCL4, CCL4L2, CCL3L3, TXNIP, CD52, CXCR4* and *CORO1A*) while activated T cells highly expressed *LTB, IL2, HSPD1, BCL2A1, HSPA1A* and *HSPA1B* (Fig. 3B; Data S3), which indicate inert T plays an important role in inflammatory responses and has weak effector functions.

**Fig. 3.**
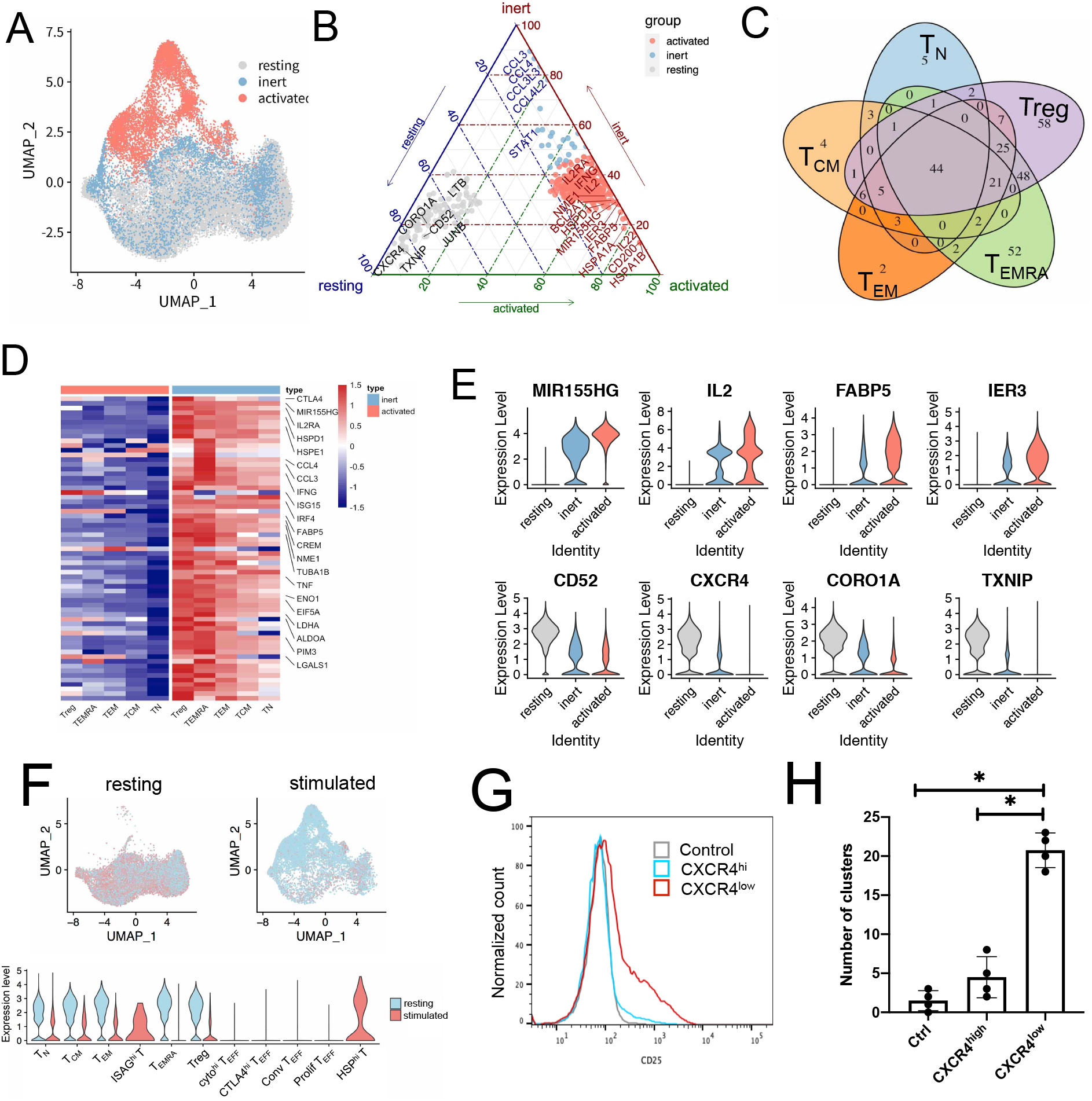
Feature of inert T and factors affecting T cell activation. **(A)** UMAP projection of CD4 T cells, colored by resting T, inert T and activated T. **(B)** Ternary diagram of T cell subsets specific genes between resting T, inert T and activated T. **(C)** Venn diagrams of inert T specific genes in T_N_, T_CM_, T_EM_, T_EMRA_ and Treg (avg_logFC >= 1). The inert T specific genes were calculated by comparing inert T to resting T in each T cell subset. **(D)** Gene expression heatmap of inert T specific genes that shared among T_N_, T_CM_, T_EM_, T_EMRA_ and Treg. **(E)** Expression level of activated T specific genes (top) and resting T specific genes (bottom) in resting T, inert T and activated T. **(F)** Expression level of *CXCR4* on UMAP projection (top), and in CD4 T cell subsets (bottom) pre-and post-stimulation. **(G)** CD25 level in control, stimulated T derived from *CXCR4*^low^ T and stimulated T derived from *CXCR4*^hi^ T. **(H)** The number of T cell aggregation or “clustering” in control, stimulated T derived from *CXCR4*^low^ T and stimulated T derived from *CXCR4*^hi^ T.

We further separated each of the five T cell subsets, namely T_N_, T_CM_, T_EM_, T_EMRA_, and Treg, into resting subset and inert subset, E.g. T_N_ was separated into resting T_N_ and inert T_N_, with differentially expressed genes between resting T_N_ and inert T_N_ similar to that between resting T and inert T (fig. S2C). The specific upregulated/downregulated genes of inert T cell subsets compared with their resting T cell counterparts are largely shared (Fig. 3C, 3D and fig. S4, D-E), indicating the features of inert T subsets are consistent among different T cell subsets. On the other hand, inert T is a specific state with many genes having intermediate expression level between resting T and activated T (Fig. 3E). *CXCR4* was moderately expressed in inert T cells, comparing its high expression in resting T cells and very low expression in activated T (Fig. 3E). Because *CXCR4* is a key CXC chemokine receptor for mediation of cell migration, high expression of *CXCR4* in resting T may facilitate the cells migration to its targeted locations (*31, 32*).

Further analyses showed expression of *CXCR4* had strong cellular heterogeneity (Fig. 3F). In particular, HSP^hi^ T highly expressed *CXCR4* while other activated T cells did not express *CXCR4*, consistent with aforementioned analyses that HSP^hi^ T are similar to T_N_ in some way (Fig. 3F). Considering the expression heterogeneity of *CXCR4*, we sorted the resting CD4 T cells into *CXCR4*^*hi*^ T and *CXCR4*^low^ T by FACS (fig. S3A). Both *CXCR4*^hi^ T and *CXCR4*^low^ T were co-stimulated by CD3 antibody and CD28 antibody for 16 hours. FACS analyses showed anti-CD3/CD28 co-stimulated CXCR4^low^ T expressed much higher level of CD25 than *CXCR4*^hi^ T (Fig. 3G and fig. S3B). Further analyses showed that the number of T cell aggregation or “clustering” in stimulated *CXCR4*^low^ T is significantly higher than that in stimulated *CXCR4*^hi^ T (Fig. 3H and fig. S3C). These results indicated that *CXCR4*^low^ T cells responded to anti-CD3/CD28 stimulation much more efficiently than *CXCR4*^hi^ T.

### Stimulated CD4 T cells showing different T cell subsets in the presence of CD8 T cells

CD3 T cells, which include CD4 T cells and CD8 T cells, is another commonly used model to study T cell activation (*33*). We performed scRNA-seq on both resting CD3 T cells and stimulated CD3 T cells. Analyses of CD3 T cells identified 9 CD4 T cell subsets and 7 CD8 T cell subsets (fig. S4A-S4E). Pseudotime analyses of CD3 T cells identified two major trajectories, which were CD4 T cell activation lineage and CD8 T cell activation lineage (fig. S4F), indicating activation of CD4 T cells and activation of CD8 T cells are two events. In this way, we could regard the CD4 T cells in stimulated CD3 T cells as stimulated CD4 T cells co-incubating with CD8 T cells (stimCo T). By integrated analyses of stimulated CD4 T cells (stim T) and stimCo T, we identified 10 CD4 T cell subsets, which essentially consist with the T cell subsets in Fig 1 (Fig. 4A; fig. S5A). A T cell subset, which highly expressed *ISG15, IFIT2, MX1, IFIT3, ISG20* and *IFI44L*, enriched GO terms such as defense response to virus (P=6.31×10^−15^) (fig. S5B-S5C), thus were called IFN^hi^ T.

**Fig. 4.**
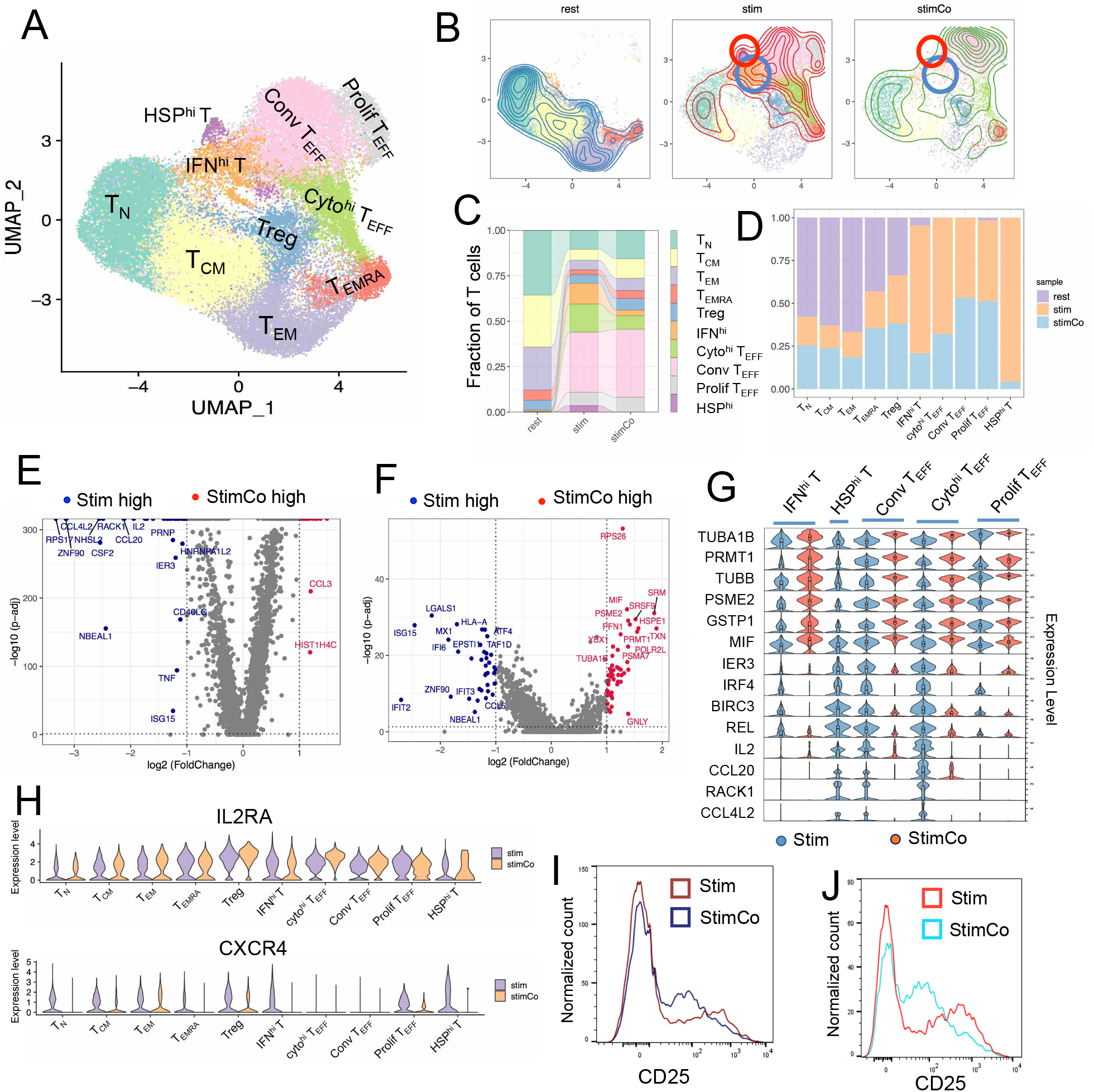
Change of stimulated CD4 T subsets and signals when stimulations co-incubated with CD8 T. **(A)** UMAP projection of CD4 T cells including rest T, stim T and stimCo T, colored by T cell subset. **(B)** Differences of cell distributions of rest T, stim T and stimCo T, with HSP^hi^ T and IFN^hi^ T marked by circles. **(C)** Fraction of T subsets in rest T, stim T, and stimCo T. The lines between two bars link the T cell subset counterparts in different samples. **(D)** Cell composition of T cell subsets based on cell sources. **(E)** DEGs between stim T and stimCo T. **(F)** DEGs of IFN^hi^ between stim T and stimCo T. **(G)** Expression level of stim T specific genes and stimCo T specific genes in activated T cell subsets. **(H)** Expression levels of *CXCR4* and *IL2RA* in T cell subsets of stim T and stimCo T. **(I)** FACS analyses of CD25 in stim T and stimCo T. **(J)** FACS analyses of CD25 in stimulated T derived from CXCR4^low^ T and stimulated T derived from CXCR4^low^ T in the presence of CD8 T.

We found that stim T and stimCo T had different distribution and different density on UMAP plot (Fig. 4B), indicating CD4 T cell activation was affected by the presence of CD8 T cells. In particular, stimCo T has higher density on conv T_EFF_ and much lower fraction of HSP^hi^ T and IFN^hi^ T compared with stim T (Fig. 4B). Quantitatively, stim T had 32.9% conv T_EFF_ and 15.6% cyto^hi^ T_EFF_, while stimCo T had 37.4% conv T_EFF_ and 7.4% cyto^hi^ T_EFF_, indicating stimCo T have increased conv T_EFF_ and decreased cyto^hi^ T_EFF_ (Fig. 4C). StimCo T has lower fraction of HSP^hi^ T (0.43%) and IFN^hi^ T (6.5%) than stim T in which HSP^hi^ was 4.92% and IFN^hi^ T was 16.1% (Fig. 4C). These results indicated stimulation of CD4 T with CD8 co-incubation might promote CD4 T differentiate into conv T_EFF_ while repress CD4 T differentiate into HSP^hi^ T, IFN^hi^ T and cyto^hi^ T_EFF_. Indeed, HSP^hi^ T, IFN^hi^ T and cyto^hi^ T_EFF_ are mainly from stim T, with few contribution from stimCo T (Fig. 4D).

### Different signals of T cell activation between stim T and stimCo T

We identified 65 differentially expressed genes (DEGs) between stim T and stimCo T. The 21 stimCo T specific genes including *TUBB, TUBA1B, MIF, GSTP1, SRM* and *HIST1H4C*, many of which involved in regulation of protein kinase and response to stimulus, indicating stimCo T have strong effector function (Fig. 4E; Data S4). The 35 stim T specific genes significantly enriched cytokine signaling (*CCL4, IL2, CCL20, CCL4L2, TNF, ISG15*) (Fig. 4E; Data S4). We further identified the DEGs between stim T and stimCo T in each T cell subset. Stim IFN^hi^ T highly expressed genes related to interleukin-mediated signaling (*MX1, OAS1, OASL, IFIT3*), while stimCo IFN^hi^ T highly expressed genes related to profilin binding, actin and tubulin folding (*ACTB, TUBA1B, DNAJA1, HSPA8*) (Fig. 4F). Stim conv T_EFF_ highly expressed genes related to cytokine signaling (*CD40LG, CSF2, IFNG, IL2, IRF1, CCL20*), while stimCo T_EFF_ highly expressed genes related to detoxification of reactive oxygen species (*GSTP1, PRDX2, TXN, CCL3, LTA, LCK*) (fig. S5D). Furthermore, cytokines or stimulation response genes (*CCL4L2, RACK1, CCL20, IL2, REL, BIRC3, IRF4*, and *IER3*) were highly expressed in stim activated T subsets (Fig. 4G). While signal transduction and cellular functional genes (*TUBA1B, PRMT1, TUBB, PSME2, GSTP1, MIF*) were highly expressed in stimCo activated T subsets (Fig 4G). Pseudotime analyses showed that trajectory of stimCo T extended and differentiated sufficiently, while trajectory of stim T terminated earlier, with a large number of cells were in the midway of the trajectory (fig. S5E-S5G). Therefore, these results support that stimulation of CD4 T cells in the presence of CD8 T cells significantly increased the immunological functions of activated T while reduced the over-response and cytokine production.

Integrated analyses showed stim T and stimCo T expressed similar level of *IL2RA* (Fig. 4H), potentially indicated stim T and stimCo T had similar activation level based on active marker. Both CD4 T cells and stimCo CD4 T cells (co-incubated with CD8 T cells) were co-stimulated by CD3 antibody and CD28 antibody for 16 hours. FACS analysis showed distribution of CD25 expression in stim T was broader and with more extreme values than that in stimCo T (Fig. 4I), indicating expression of CD25 in stimCO T is more homogenous than stim T. On the other hand, most stimCo T cell subsets expressed *CXCR4* significantly lower than stim T cell subsets (Fig. 4H), indicating that stimCo T responds to stimulation more efficiently than stim T by function. We further stimulated *CXCR4*^low^ T and analyzed the expression of CD25 by FACS. Distribution of CD25 expression in stimulated T derived from *CXCR4*^low^ T was much broader than their counterpart in the presence of CD8 T cells (Fig. 4J), supporting the presence of CD8 T made the response of CD4 T become more homogeneous. These results potentially indicate that CD8 T could significantly change the stimulated T cell subsets and expression signals by cell-cell communication or its secreting cytokines.

### The effects of different factors on CD4 T cell activation

To explore the factors affecting CD4+ T cell activation, we further integrated scRNA-seq data of naïve CD4+ T cells (restT_N_), stimulated CD4 T cells derived from CD4 T_N_ (T_N_stim) and T_N_stim in the presence of different cytokine cocktails (*20*). The integrated 5 data sets are restT_N_, T_N_stim, T_N_stim in the presence of IL2 and TGFβ (T_N_stim+IL2), T_N_stim in the presence of IL4 and anti-IFNγ (T_N_stim+IL4), T_N_stim in the presence of IL6, IL23, IL1β, TGFβ, anti-IFNγ and anti-IL4 (T_N_stim+IL6). The integrated 60,639 CD4 cells were re-clustered into 11 cell subsets, including 5 resting T cell subsets (T_N_ #1, T_N_ #2, T_CM_, T_EM_ and T_EMRA_) and 6 activated T cell subsets (IFN^hi^ T, T_EFF_, Th0, prolif T #1, prolif T #2, and HSP^hi^ T) (Fig. 5A and fig. S6A). The cell composition of the 8 samples displayed three distinct patterns: rest pattern representing resting T cell (rest and restT_N_), T_N_stim pattern representing stimulated T derived from T_N_ (T_N_stim, T_N_stim+IL2, T_N_stim+IL4, T_N_stim+IL6) and stim pattern representing stimulated T derived from peripheral CD4 T (stim and stimCo) (Fig. 5B, 5C). Majority of the cells in rest pattern are T_N_ #1 and T_N_ #2, which is quite different from stimulated T cells including stim pattern and stimT_N_ pattern (Fig. 5B, 5C). Stim pattern (stim and stimCo) have the highest fraction of T_EFF_ (Fig. 5C). Compared with stim pattern, T_N_stim pattern has increased Prolif T, ISAG^hi^ T, HSP^hi^ T and Th0. The slight differences among the samples in T_N_stim pattern compared with their differences with stim pattern, suggesting that cytokines have weaker effect on T cell activation compared with type of resting T cells (Fig. 5C).

**Fig. 5.**
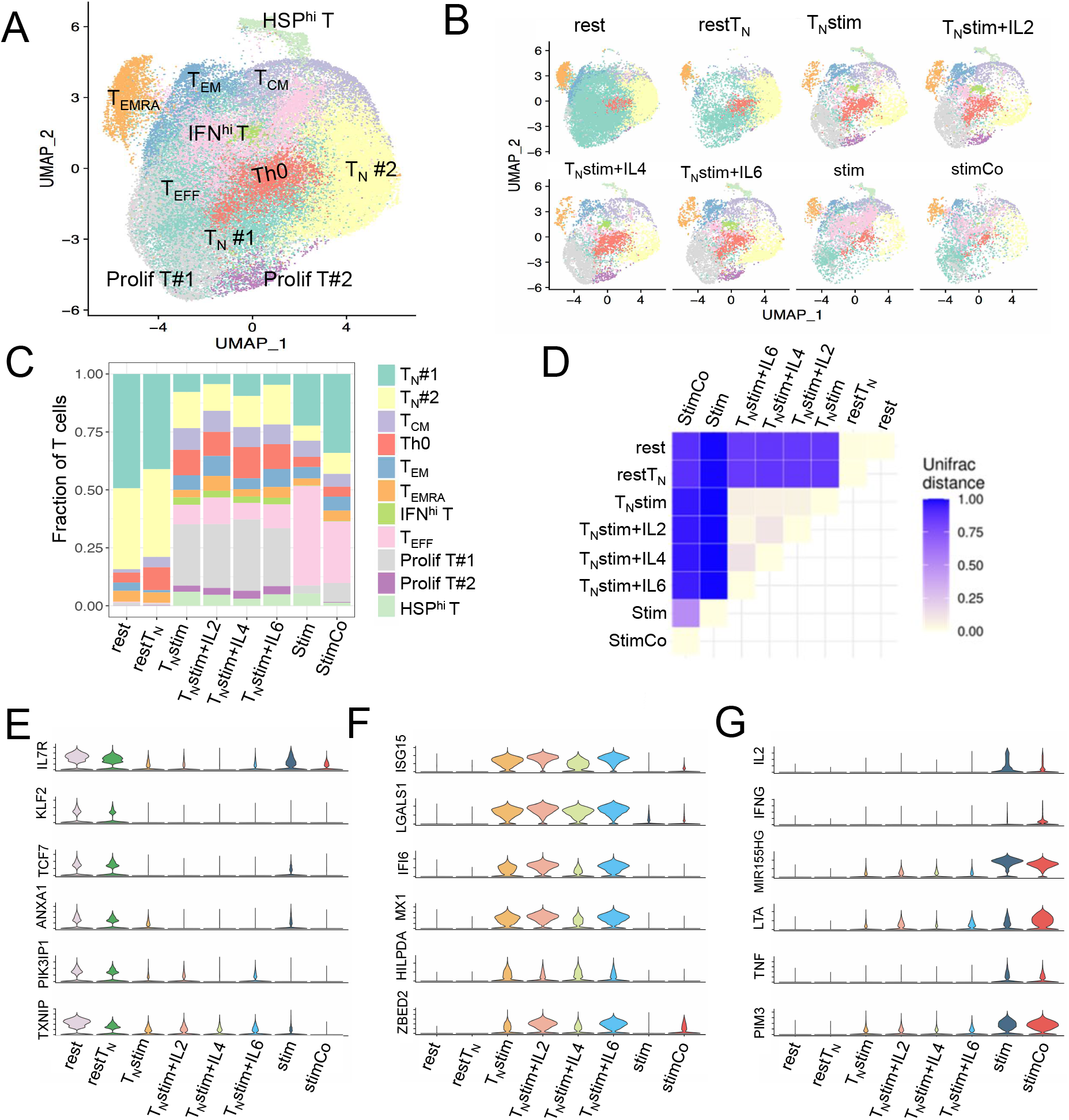
Exploring multiple factors affecting CD4 T cell activation by integrating 8 samples. **(A)** UMAP projection of integrated CD4 T cells from 8 conditions, colored by cell subset. **(B)** UMAP projection of CD4 T cells in each condition, namely, rest, restT_N_, T_N_stim, T_N_Stim+IL2, T_N_Stim+IL4, T_N_Stim+IL6, stim and stimCo. **(C)** Bar plot of fraction of T cell subsets in each condition. **(D)** UniFrac distances between CD4 T cell samples summarized in a correlation plot. The values of UniFrac distance range from 0 to 1, with lower value indicating higher similarity. **(E-G)** Expression levels of rest pattern specific genes **(E)**, T_N_stim pattern specific genes **(F)** and stim pattern specific genes **(G)** in each sample.

We further calculated UniFrac distance (*34*) to quantify the similarity of cell composition between two samples. UniFrac distance ranges from 0 to 1, where 0 indicates two samples have the same cell composition and 1 indicates two samples have entirely separate clusters. UniFrac distances between samples within each of the rest pattern, T_N_stim pattern and stim pattern are very small, supporting that the internal pattern differences is much weak. Meanwhile, uniFrac distances between samples across patterns are much higher than that within pattern, indicating that type of resting T cells is the dominant factors affecting T cell activation (Fig. 5D). While the UniFrac distance between stim and stimCo is the highest among all within pattern comparisons, indicating co-incubation of the CD8 T is second strongest factor affecting T cell activation (Fig 5D). Indeed, stimCo T has quite different distribution on UMAP comparing with the other stim or T_N_stim (fig. S6B). Samples in rest pattern highly expressed resting T cell specific genes such as *TCF7, KLF2, ANXA1, TXNIP*, and *IL7RA* (Fig. 5E). Samples in T_N_stim pattern (T_N_Stim, T_N_stim+IL2, T_N_stim+IL4 and T_N_stim+IL6) highly expressed interferon associated genes such as *ISG15, MX1* and *IFI6*, (Fig. 5F), consistent with our observation that these samples have the highest fraction of IFN^hi^ T. Samples in T_N_Stim pattern also highly expressed *LGALS1, ZBED2* and *HILPDA* that promote cell proliferation (*35*), consistent with pronounced increase of proliferation T. Samples in stim pattern (stim and stimCo) highly expressed *PIM3, TNF, LTA, MIR155HG, IFNG* and *IL2* (Fig. 5G), consistent with the pronounced increase of T_EFF_ in these samples.

## Discussion

CD4 T cells are one of the most commonly used models for investigation of T cell activation. Most of the previous studies focused on naïve CD4 T cells because they are relatively homogeneous (*4*) and can differentiate into various functional T cell subsets (*8, 20*). In order to analyze the factors affecting T cell activation, we selected the most easily obtainable CD4 T cells, peripheral CD4 T cells, to initiate the study. Although the peripheral CD4 T cells is highly heterogeneous, including CD4 T_N_, CD4 T_CM_, CD4 T_EM_, CD4 T_EMRA_ and ISAG^hi^ T (*4, 33*), we identified the signals of T cell activation in the heterogeneous sample due to the analyses was conducted at single cell level. Among the 4 T_EFF_ subsets generated after stimulation, cyto^hi^ T_EFF_ expressed the highest level of *IL2* and *IFNG*, indicating cyto^hi^ T_EFF_ are mainly Th1. However, we did not separate the CD4 T_EFF_ into different T helper subsets because we focused on the genome-wide expression profile. Among the stimulated CD4 T cells, HSP^hi^ T enriched HSP genes and genes related to cellular responses to stress. HSP^hi^ T were mainly projected on T_N_ when we re-performed UMAP projection using genes without HSP genes, indicating HSP^hi^ T have an expression profile similar to T_N_ except highly expressed HSP genes. We also identified a stimulated T cell subset that highly expressed interferon associated genes, namely IFN^hi^ T. Overall, majority of resting CD4 T cells developed into various effector T cells upon anti-CD3/CD28 stimulation, while a small fraction of CD4 T cells develop into specific status such as HSP^hi^ T and IFN^hi^ T.

The existence of inert T indicates some resting T cells respond to stimulation much slower than the other T cells due to the heterogeneous of resting CD4 T cells. We found that inert T subsets specific genes compared with their resting T counterparts are shared, indicating shared mechanisms of T cell activation between inert T subsets. Inert T highly expressed activated T specific genes (*IL2RA, NME1, MIR155HG, FABP5, IL2, CD200* and *IER3*), indicating inert T is not complete quiescent and have responded to the anti-C3/CD28 stimulation. On the other hand, Inert T cells highly expressed cytokine related genes (*CCL3, CCL4, CCL4L2, CCL3L3, TXNIP, CD52, CXCR4* and *CORO1A*) compared with activated T and resting T, indicating inert T plays important role in inflammatory responses and have weak effector functions. Furthermore, many genes were intermediately expressed in inert T, indicating that inert T is a transitional state between resting T and activated T. The expression level of *CXCR4* decreased successively in resting T, inert T and activated T, with expression level showed high heterogeneity among T cell subsets. We found that *CXCR4*^low^ T respond to CD3 antibody and CD28 antibody stimulation more efficiently than *CXCR4*^hi^ T, indicating the expression level of *CXCR4* significantly affecting T cell activation.

Comparison of stimulated T cell under several conditions showed that the type of resting T cells has the strongest effects on T cell activation, stimulation of peripheral CD4 T and CD4 T_N_ resulting in quite different stimulated T cell subsets. In particular, stimulated T derived from peripheral CD4 T has much higher fraction of T_EFF_ while stimulated T derived from CD4 T_N_ has much higher fraction of prolif T. The second important factors affecting T cell activation is whether the stimulation was co-incubated with CD8 T cells. Our analyzed showed CD4 T cells were more likely to differentiate into T_EFF_ and were less likely to differentiate into HSP^hi^ T and IFN^hi^ in the presence of CD8+ T cells. The effects of CD8 T cells co-incubation on T cell activation maybe mediated by cell-cell communications and/or specific microenvironment shaped by their secreted cytokines. Finally, presence of cytokine cocktail during stimulation also shaped the T cell subsets in some way, but do not have strong effects based on their expression profiles. In short, this study facilitates our understanding of the dynamics of T cell subsets upon stimulation, signals and factors affecting T cells activation.

## Materials and Methods

### Isolation of peripheral blood mononuclear cells (PBMCs)

This study was approved by IRB at Southern University of Science and Technology (SUSTech). The human peripheral blood samples used in this study were obtained from two healthy adult donors. All experiments were conducted following the protocols approved by IRB at SUSTech. PBMCs were isolated by density-gradient sedimentation of peripheral blood using Lymphoprep™ (Axis-Shield, Oslo, Norway) following previous study (*4*). Briefly, peripheral blood sample was diluted with an equal volume of PBS containing 2% FBS, transferred into a Lymphoprep™ tube. Following centrifuged at 800 × g for 20 min at room temperature, the middle white phase was transferred to a fresh tube. The enriched mononuclear cells were washed with PBS containing 2% FBS and centrifuged at 500 × g for 5 min twice.

### Isolation of CD3 T cells and CD4+ T cells

The CD3+ cells and CD4 T cells were enriched from PBMCs using CD3 MicroBeads (Miltenyi biotec, #130-050-101, German) and CD4 MicroBeads (Miltenyi biotec #130-045-101, German), respectively, following products’ instructions. In brief, PBMCs were re-suspended and washed with PBS buffer, and magnetically labeled with CD3 MicroBeads or CD4 MicroBeads. Then the cell suspension was loaded onto a MACS® Column in the magnetic field of a MACS Separator. The magnetically labeled CD3+ cells or CD4+ T cells were retained on the column. After removal of the column from the magnetic field, the magnetically retained CD3+ cells or CD4+ cells were collected by elution.

### T cell activation by anti-CD3/CD28 stimulation

MACS enriched CD4+ T cells and CD3+ T cells were stimulated with Dynabeads™ Human T-Activator CD3/CD28 for T Cell Expansion and Activation (ThermoFisher, Cat# 11131D, USA) following the manufacturer’s instructions. Briefly, 8 × 10^4^ CD4+ T cells were suspended with 100–200 µL medium. Pre-washed Dynabeads® was added at the bead-to-cell ratio of 1:1 and incubated in a humidified CO_2_ incubator at 37°C. After 16 hours, stimulated CD4+ T cells were harvested by removal of Dynabeads®, and then stained with anti-CD25 (Cat #12-0259-42, eBioscience) for FACS analyses of the efficiency of T cell activation. The CD3 T+ cells were stimulated with Dynabeads™ Human T-Activator CD3/CD28 for T Cell Expansion and Activation following exactly the same protocol as stimulation of CD4+ T cells.

### scRNA-seq library preparation and sequencing

MACS enriched CD4+ T cells (resting CD4 T cells) and anti-CD3/CD28 stimulated CD4+ T cells (stimulated CD4 T cells) were directly processed for scRNA-seq with Chromium™ Single Cell 3’ Library & Gel Bead Kit v2 (10x Genomics, Pleasanton, CA) following manufacturer’s instructions. For each sample, cells were counted in a haemocytometer chamber and about 1.6 × 10^4^ T cells were loaded into single inlet of a 10x Genomics Chromium controller to generate the nanoliter-sized Gel beads in Emulsions (GEMs). After GEM collection, cell lysis, RNA-capture and barcoded reverse transcription were performed inside each GEM, thus mRNA was reverse transcribed into cDNA. After breaking the oil droplets, libraries were generated from the cDNAs with Chromium Single Cell 3′ Library Construction Kit v2 (PN-120237, 10x Genomics). Sequencing libraries were loaded at 2.4 pM into Illumina NovaSeq 6000 with 2×150 paired-end kits using the following read length: 28 bp Read1, 8 bp I7 Index, 8 bp I5 Index and 91 bp Read2.

### Integration of public data and data pre-processing

In order to systematically analyze the factors affecting T cell activation, we integrated public scRNA-seq datasets including scRNA-seq data of Naïve CD4 T cells and diversified anti-CD3/anti-CD28 stimulated CD4 T cells from Cano-Gamez (*20*), scRNA-seq data of resting CD4 T cells (resting_2) and anti-CD3/CD28 stimulated CD4+ T cells (stimulated_2) from Ding (*36*), and scRNA-seq data of resting CD3 T cells (CD3 resting_2) from Massoni-Badosa (*37*).

We mapped reads to reference genome (hg19) and generated single-cell gene expression matrix for each scRNA-seq dataset using Cell Ranger (v3.0). We used Seurat (*38*) for quality control and basic pre-processing following our previous studies (*4, 39, 40*). We filtered out the cells with the following criteria: 1) number of detected genes <200; 2) UMI counts >40,000; 3) mitochondrial percentage >7.5% (*41*), 4) percentage of CD8 and erythrocyte gene >0.01%.

### Dimension reduction and visualization of scRNA-seq data

Data integration, data normalization, dimension reduction and visualization were performed using Seurat (*38*), similar to our previous studies (*4, 39, 40*). We identified the top 3,000 high variable genes using *FindVariableFeatures* function and further scaled the gene expression matrix using *ScaleData* function. We conducted principal component analysis (PCA) for linear dimension reduction using *RunPCA* function and the top 30 PCs were used for further analysis. We visualized cells using Uniform Manifold Approximation and Projection (UMAP) (*42*). Clusters were divided by an unsupervised graph-based clustering method.

### Identification of cell subsets and differential gene analysis

The cell clusters were annotated using T cell subset specific gene markers from reference articles. In order to identify the differentially expressed genes between two subsets or two conditions, we utilized function FindMarkers to exert differential expression analysis on cluster pairs. It returned average log Fold Change (avg_logFC) and adjusted p value (p_val_adj) for each gene. The genes with p_val_adj <0.05 and avg_logFC>1 were kept as the differentially expressed genes (DEGs).

### Gene Ontology (GO) enrichment analysis

In order to better understand the functions of cluster specific genes or DEGs between two T cell subsets, GO enrichment analyses were conducted on these gene lists using Metascape (http://metascape.org) (*43*).

### Pseudotime inference

Monocle3 (*44*) uses pseudotime to measures how a cell moves through biological progress. We used monocle3 to infer trajectories of T cell activation on UMAP space and identified trajectory-associated differential genes. Naïve T cells were selected as root node manually based on biological knowledge. The top 10 trajectory lineage associated genes were selected by the morans_I index.

### Comparison of CXCR4^hi^ T and CXCR4 ^low^ T on T cell activation

In order to analyze whether *CXCR4* impact on T cell activation, CD4+ T cells or CD3+ T cells were sorted into *CXCR4*^hi^ T cells and *CXCR4*^low^ T with anti-CXCR4 (Cat # 306510, Biolegend) on FACS Aria-III (Becton Dickinson). FACS sorted *CXCR4*^hi^ T cells and *CXCR4*^low^ T cells were co-stimulated with anti-CD3 antibody and anti-CD28 antibody. Briefly, 8 × 10^4^ FACS sorted *CXCR4*^hi^ T cells or *CXCR4*^low^ T cells were suspended with 100–200 µL medium, respectively. The anti-CD3 and anti-CD28 was added at the antibody-to-cell ratio of 1:1 and incubated in a humidified CO_2_ incubator at 37°C. After 16 hours, activated T cells were harvested and stained with anti-CD25 (Cat # 12-0259-42, eBioscience) for FACS analyses of the efficiency of T cell activation. FlowJo (version 10) was used for FACS data analyses and Fig. plot.

## Acknowledgments

We thank all members from the Jin lab for the helpful discussion. We acknowledge the assistance of Core Research Facilities of SUSTech. We thank Xibin Lu for the excellent support of FACS. The computational work was supported by Center for Computational Science and Engineering at SUSTech.

## Author contributions

W.J conceived the project. X.W collected peripheral blood. H.L conducted T cell stimulation and FACS analyses. Q.Z conducted single cell RNA-seq. H.L and Y.L analyzed the data with help from X.S and S.Y. W.J and N.H supervised the project. H.L. and W.J. prepared the manuscript, with all authors’ contribution.

## Data and materials availability

The sequencing data has been deposited in Genome Sequence Archive in BIG Data Center under accession number HRA002777.

