## Supplementary Materials for "Dynamics, signals and influencing factors of CD4 T cell activation revealed by single cell RNA-seq"

**This PDF file includes:**

Figs. S1 to S6

**Other Supplementary Materials for this manuscript include the following:**

Data S1 to S4

Fig. S1

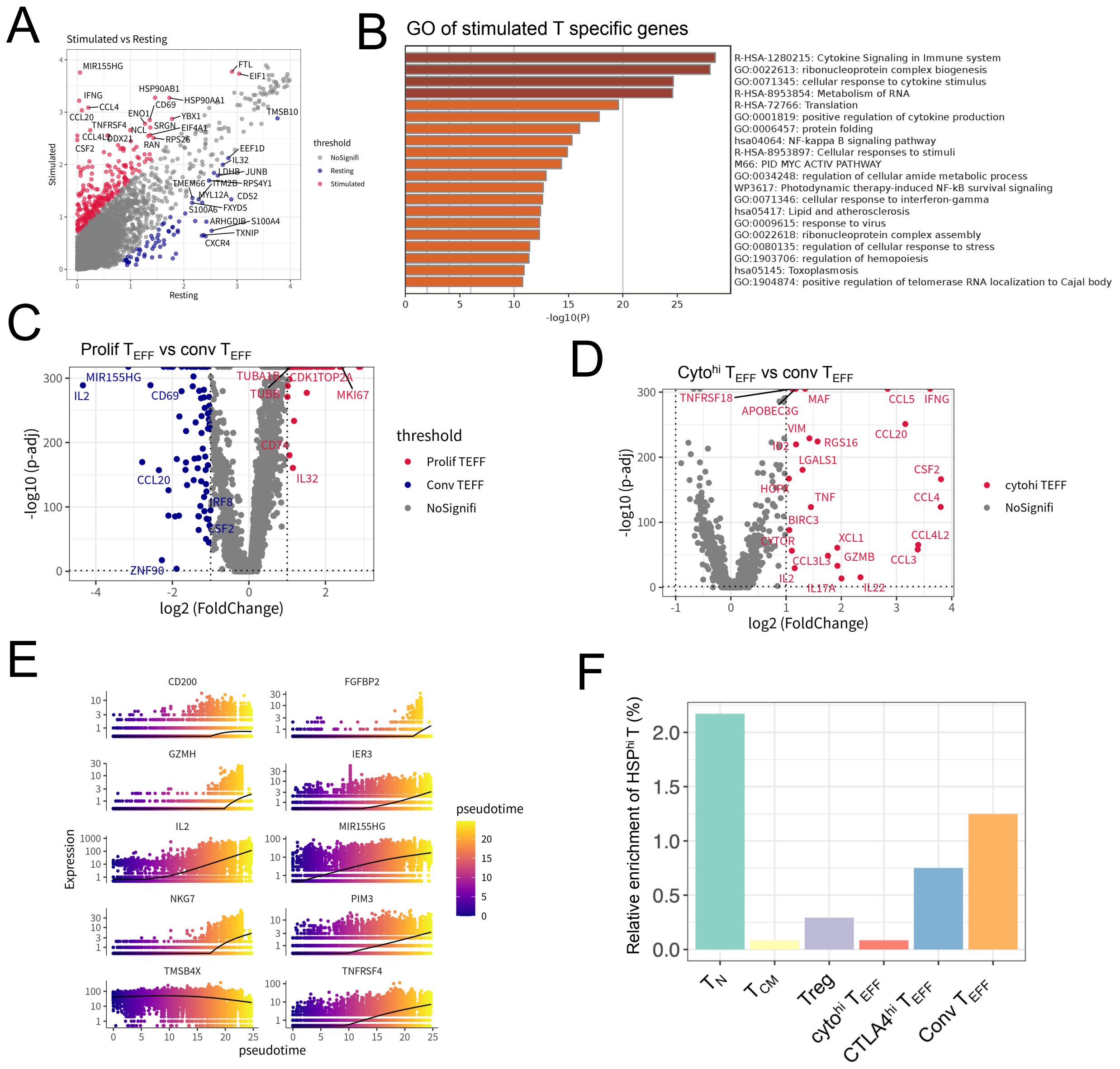


**Fig. S1. Feature and subsets of CD4 T pre-and post-anti-CD3/CD28 stimulation. (A)** Scatter plots of gene expression level in resting CD4+ T cells and stimulated CD4+ T cells. Blue dot and red dot represents resting T cell specific genes and stimulated T cell specific genes, respectively. **(B)** GO enrichment analysis of stimulated CD4 T cell specific genes. **(C)** Volcano plot of DEGs between conv $T_{EFF}$ and prolif $T_{EFF}$. Red points represent prolif $T_{EFF}$ specific expressed genes, while blue points represent conv $T_{EFF}$ specific expressed genes. **(D)** Volcano plot of DEGs between conv $T_{EFF}$ and $cyto^{hi}$ $T_{EFF}$. Red points represent $cyto^{hi}$ $T_{EFF}$ specific expressed genes. **(E)** Genes associated with T cell activation lineage. **(F)** Bar plot of number of $HSP^{hi}$ T normalized to its projected T cell subsets.

Fig. S2

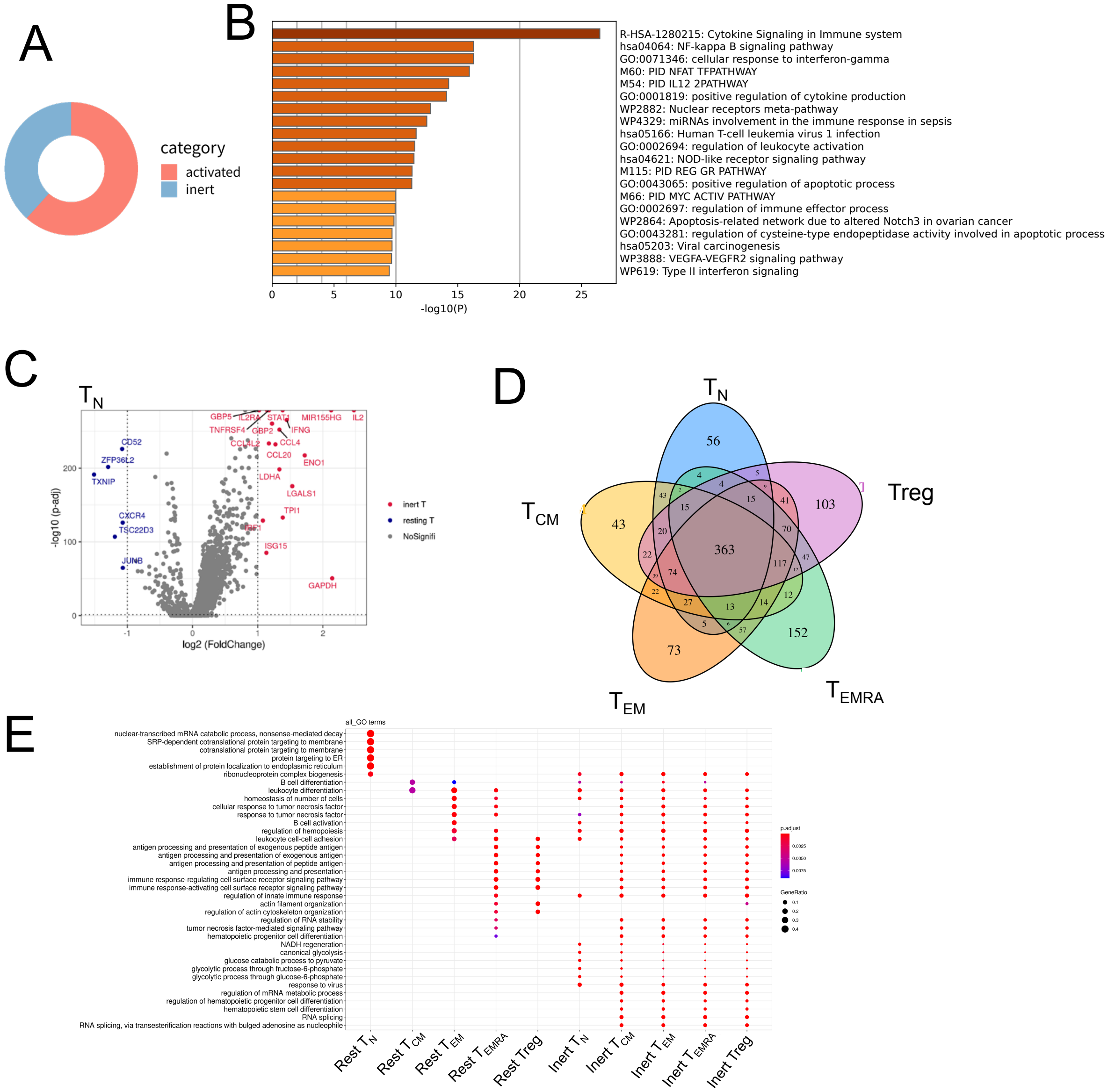


**Fig. S2. Feature and subsets of inert T. (A)** The proportion of inert T and activated T cells to the total stimulated T cells. **(B)** GO analysis of inert T subset specific genes compared with resting T cells. **(C)** Volcano plot of DEGs between inert $T_N$ and resting $T_N$. Red points represent inert $T_N$ specific expressed genes, while blue points represent resting $T_N$ specific expressed genes. **(D)** Venn diagrams showing majority of inert T subset specific genes shared among $T_N$, $T_{CM}$, $T_{EM}$, $T_{EMRA}$ and Treg (avg_logFC >= 0.25). The inert T subset specific genes were calculated by comparing inert T to resting T in each T cell subset. **(E)** GO analysis of inert T subset specific genes and resting T subset specific genes (avg_logFC >= 0.25) showing the enriched GO terms of inert T subset specific genes were shared. The color of the dot represented the adjustive p value, and the size of the dot represent-ed the ratio of expressing genes counts to all genes counts.

Fig. S3

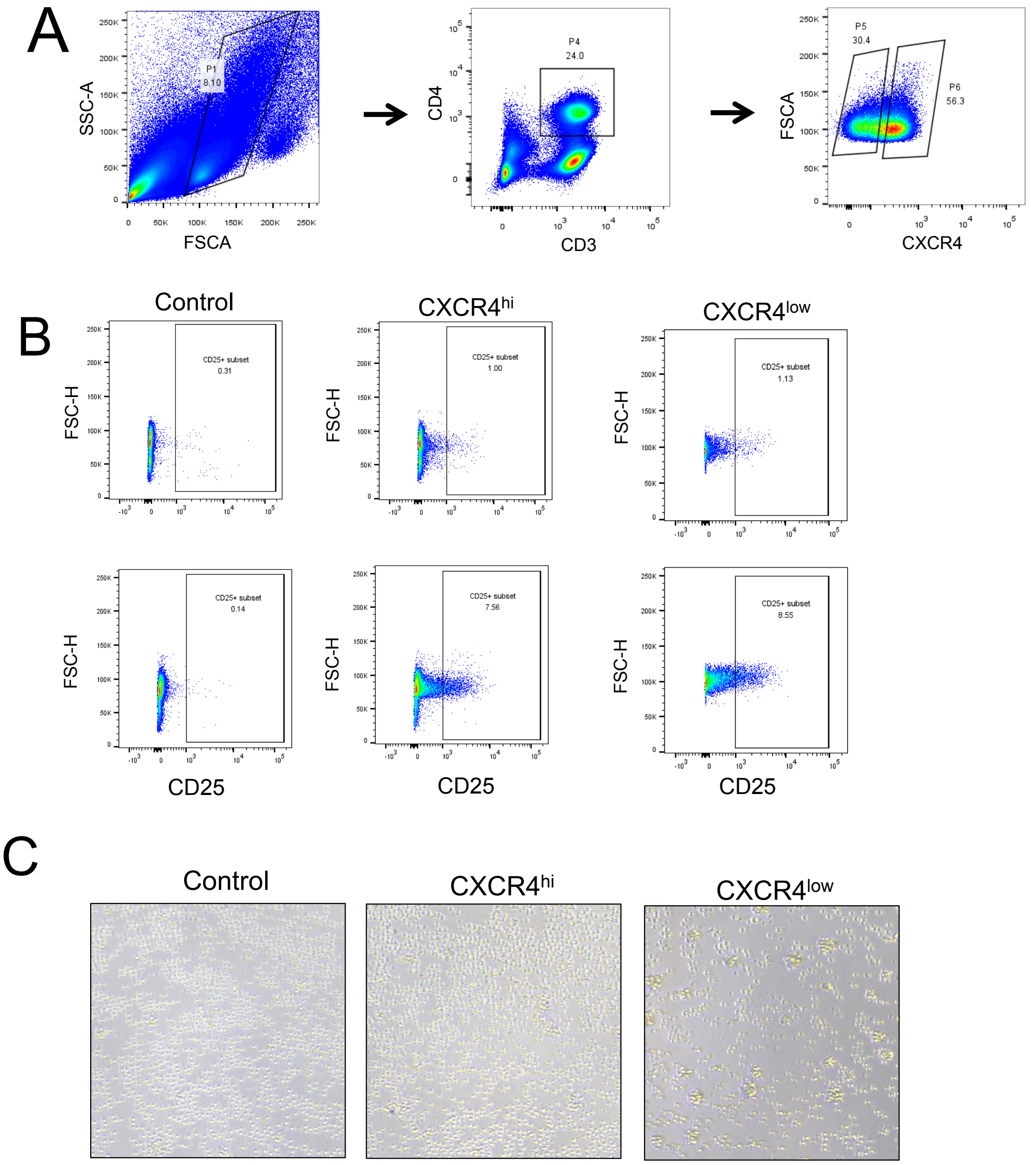


**Fig. S3. Comparisons of T cell activation between *CXCR4*$^{high}$ CD4+ T cells and *CXCR4*$^{low}$ CD4 T cells. (A)** Sorting of CXCR4$^{hi}$ CD4 T cells and CXCR4$^{low}$ C4+ T cells from PBMCs by FACS. **(B)** CD25 level in control (left), anti-CD3/CD28 stimulated CXCR4$^{low}$ CD4 T cells (middle) and anti-CD3/CD28 stimulated CXCR4$^{hi}$ CD4 T cells by FACS sorting (right). **(C)** Images of T cell aggregation or "clustering" in control, anti-CD3/CD28 stimulated CXCR4$^{low}$ CD4 T cells and anti-CD3/CD28 stimulated CXCR4$^{hi}$ CD4 T cells.

Fig. S4

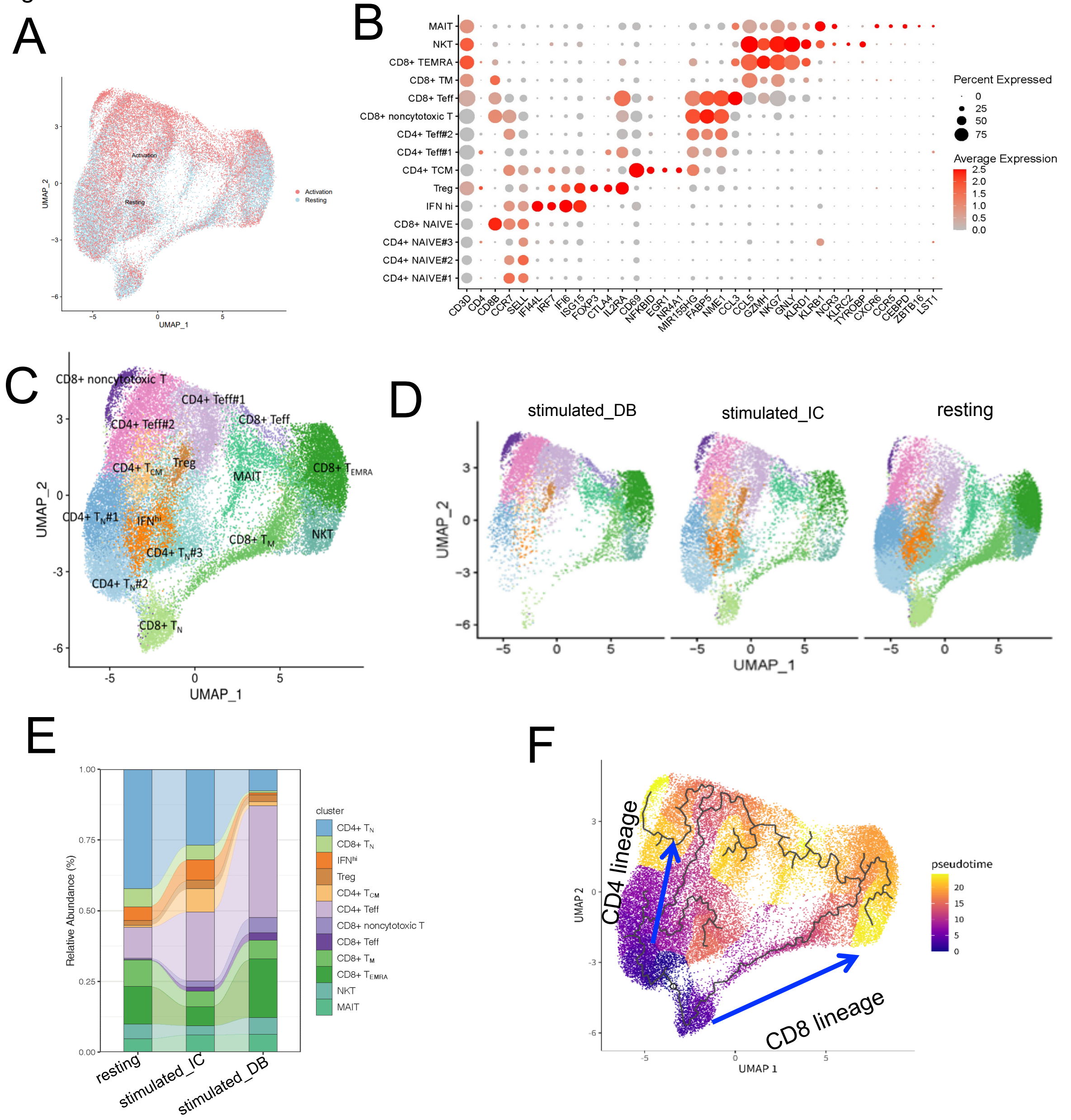


**Fig. S4. T cell subsets and lineage of CD3 T cell activation. (A)** UMAP projection of CD3 T cells pre-and post-stimulation, colored by whether it was stimulated. **(B)** Normalized expression level and expression percentage of the cell-type-specific genes in the 15 T cell subsets. Color represents normalized expression level, and dot size represents expression percentage. **(C)** UMAP projection of CD3+ T cells pre-and post-stimulation, colored by T cell subset. **(D)** UMAP Projection of resting T cells, stimulated T cells by ImmunoCult™ (stimulated_IC) and stimulated T cells by Dynabeads™ (stimulated_DB), colored by cell subset. **(E)** Fraction of T cell subsets in resting T cells, stimulated_IC, and stimulated_DB. The lines between two bars link the T cell subset counterparts in different samples. **(F)** Two distinct lineages, namely CD4+ T lineage and CD8+ T lineage, were displayed in CD3 T cell activation process.

Fig. S5

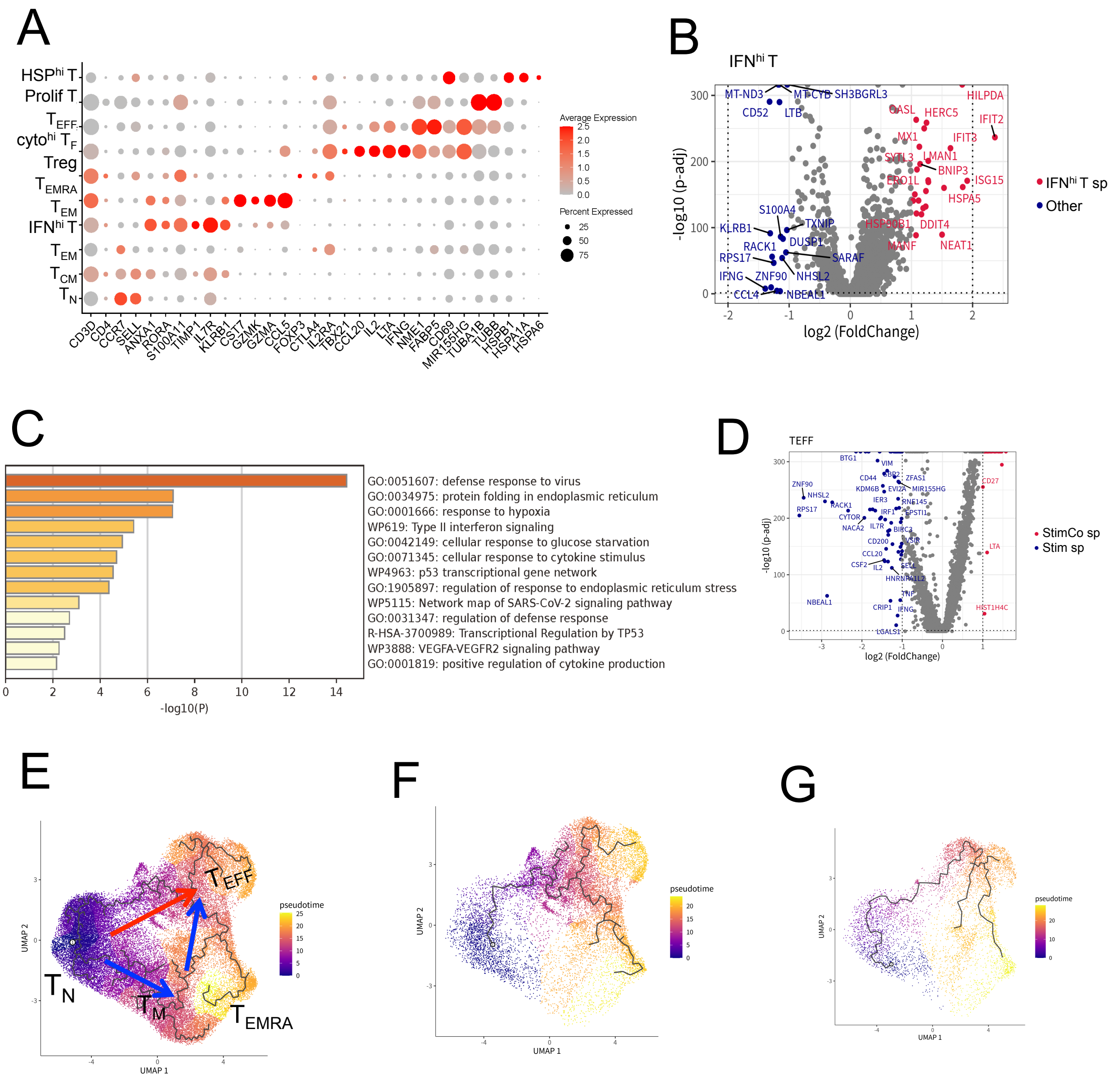


**Fig. S5. Integrated stimulated CD4 T cells co-incubated with CD8 T cells. (A)** Normalized expression level and expression percentage of cell-type-specific genes in the 10 subsets. **(B)** Volcano plot of DEGs between $IFN^{hi}$ T and other T cells. **(C)** GO analysis of $IFN^{hi}$ T specific genes. **(D)** Volcano plot of DEGs between stimCo $T_{EFF}$ and $T_{EFF}$. **(E)** Inferred two main lineages in stimulated CD4+ T cells: $T_N$->$T_{EFF}$; $T_N$->$T_M$->$T_{EFF}$. **(F)** Inferred lineage in stim T. **(G)** Inferred lineage in stimCo T.

Fig. S6

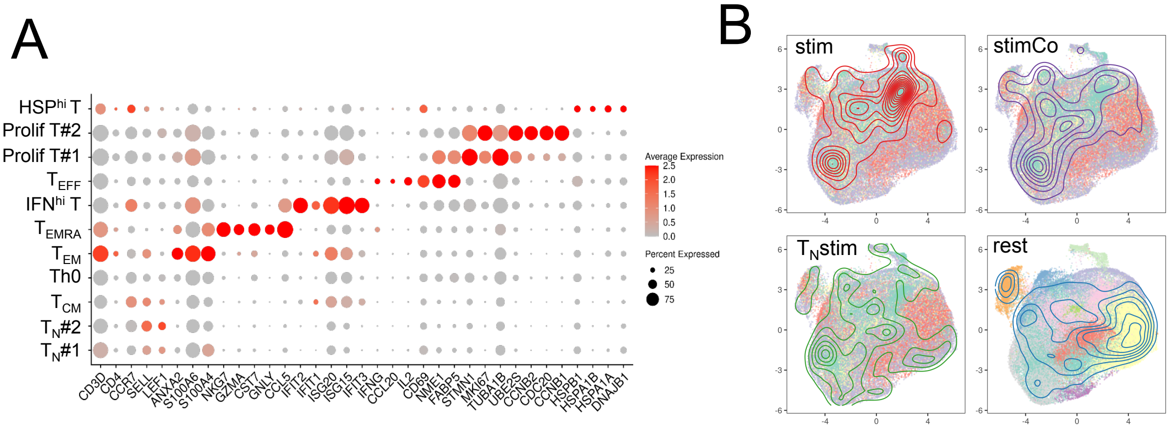


**Fig. S6. Analyses of CD4 T cell activation with multiple samples. (A)** Normalized expression level and expression percentage of cell-type-specific genes of the 11 subsets in integrated T cells. **(B)** UMAP projection of the CD4 T cells with 4 major conditions, namely rest, stim, stimCo, and $T_N$stim.

**Data S1. HSP$^{hi}$ T specific genes comparing with other T cells.**

**Data S2. DEGs between inert T and resting T.**

**Data S3. DEGs between inert T and activated T.**

**Data S4. DEGs between stimT and stimCo T.**
